# Managed culls mean extinction for a marine mammal population when combined with extreme climate impacts

**DOI:** 10.1101/2021.09.29.462338

**Authors:** Katrina J Davis

## Abstract

Human actions led to the worldwide decline of marine mammal populations in the 18^th^–19^th^ centuries. However, the global uptake in protective legislation during the 20^th^ century has recently allowed many marine mammal populations to recover. This positive trend is particularly true of pinnipeds (e.g., seals and sea lions), whose recovering populations are increasingly in conflict with fisheries. Many fisheries organisations call for managed culls of sea lion populations to reduce competition for target fish species as well as damage to catch and fishing gear through operational interactions. However, despite widespread perceptions that sea lion populations are generally increasing, to-date culls have been considered or implemented without quantitative evidence of their impacts on seal lion population viability. This knowledge gap is particularly concerning given the expected increase in extreme climate conditions, such as extreme El Niño events, which together with culls could push sea lion populations in some parts of the world into the extinction vortex. In this analysis, I develop and parameterise stochastic matrix population models of the South American sea lion (*Otaria flavescens*) to project the impact of (1) three cull scenarios with different intensity and temporal frequency targeting adult females, (2) extreme El Niño events whose frequency is modelled using a Markovian transition matrix, (3) and the interaction of culls and extreme climate events on population dynamics. I focus on the Chilean population of *O. flavescens*, where recent increases in sea lion numbers have triggered widespread conflict with small-scale fisheries, and where sea lion populations will increasingly be affected by extreme El Niño conditions. I find that sea lion populations decline below minimum viable population sizes under all scenarios involving culls and extreme climate events. By explicitly considering parameter uncertainty, this approach is a call to action for future research to focus on collecting stage-specific, annual population data to reduce uncertainty regarding marine mammal vital rates.

## Introduction

Human actions, notably hunting, led to the worldwide decline of marine mammal populations in the 18^th^—19^th^ centuries (Gerber and Hilborn, 2001). Fortunately, the global uptake in protective legislation during the 20^th^ century has allowed many populations to bounce back (Magera et al., 2013). This is particularly true of pinnipeds, including sea lions (Milano et al., 2020), whose recovering populations are now increasingly in conflict with fisheries (Cook et al., 2015; Scordino, 2010). The exact economic impacts of conflict between fisheries and sea lion populations are unknown. Regardless, the perception that these impacts are large and increasing has led many fisheries organisations to call for managed culls of sea lion populations. The expectation is that reducing sea lion populations will reduce competition for target species (*i.e*., biological or predatory interactions (Beverton, 1985)), and damage to catch and fishing gear (*i.e*., operational interactions (Beverton, 1985)). However, before managers can sanction population culls, it is essential to understand the projected impacts of these culls on the viability of natural populations. For a population to be viable, it must be able to withstand stochastic perturbations, e.g., environmental stochasticity or natural catastrophes, in the long-term given its specific biogeographic setting (Shaffer, 1981). Thus, a robust population viability assessment of sea lions must incorporate the impact of future climate change, which may have profound consequences for these species (de Oliveira et al., 2012; Kovacs et al., 2012).

To-date, most pinniped demographic research has had a methodological focus—with the objective of developing population models when demographic data is limited. This includes work by Kauhala et al. (2012), who estimated demographic structure and mortality rates of the Baltic grey seal population based on the age structure present in hunted grey seals (*Halichoerus grypus*), and Wielgus et al. (2008) who used inverse methods to estimate demographic rates and asymptotic population growth rates for the California sea lion (*Zalophus californianus*). Other analyses have focused on assessing the impact of site selection and data aggregation decisions on estimates of pop survival in grey seals (Engbo et al., 2020). By contrast, the contribution of the current analysis is to assess the impact of disturbances on population viability—specifically, managed culls and extreme climate events—as well as to explicitly incorporate uncertainty regarding vital rates into the model architecture. This analysis therefore provides a first step towards understanding the potential impact of managed culls and extreme climate events on the population dynamics of a marine mammal species under high uncertainty regarding model parameters.

Here, I overcome limitations in past approaches by building a stochastic demographic model that incorporates parameter uncertainty in a species that is at the center of much human-wildlife conflict: the South American sea lion (*Otaria flavescens*, (Cappozzo and Perrin, 2009)). Specifically, I explore the potential impacts of managed culls and projected climate change impacts on the population dynamics of *O. flavescens* in Chile. Recovery of this species (*O. flavescens*) has led to widespread conflict with small-scale fisheries (Davis et al., 2021), particularly along the coast of Chile. Although Chile does not currently cull sea lion populations, a moratorium on sea lion harvest is about to expire in 2021 (Decreto Exento N° 31-2016), thus creating a legal loophole for which quantitative evidence is urgently missing. Culls of sea lion populations are widely demanded by fishers in this region (Davis et al., 2021), and hence there is a need to assess their potential impacts on the species. However, this region is also affected by extreme El Niño conditions, which have previously had large negative impacts on pinniped populations in the north of the Chile and also in neighbouring Peru (de Oliveira et al., 2012; Sepúlveda et al., 2015). Extreme el Niño events are characterised by sea surface temperatures exceeding 28°C (Cai et al., 2014). These conditions lead to large decreases in prey availability—causing decreases in fecundity and increases in juvenile and adult sea lion mortality (de Oliveira et al., 2012).

To assess the impact of managed culls and extreme climate events on the Chilean population of *O. flavescens*, I combine known and imputed vital rates (*i.e*., stage-specific survival, development, and reproduction) for *O. flavescens* to parameterise stochastic matrix population models. I then project the *O. flavescens* population under a range of scenarios and impacts. Scenarios include a base case scenario—absent of any cull or extreme climate impacts, as well as three cull scenarios with different intensity and temporal frequency targeting adult females, and an extreme climate scenario where extreme El Niño conditions impact sea lion vital rates at frequencies expected for the coming century. Finally, I assess the combined impact of culls and extreme climate events on sea lion population size through time. I hypothesise that the impact of managed culls or extreme climate events in isolation will not adversely affect the viability of this population. However, I expect that the population will decrease below a minimum viable population size, thus become functionally extinct (*sensu* Shaffer, 1981), under managed culls in combination with projected frequencies of extreme El Niño conditions. I find that the Chilean sea lion population declines below a minimum viable population size under all scenarios involving culls and extreme climate events. Based on these findings, I provide quantitatively-based recommendations for the management of human-wildlife conflict between pinnipeds and fisheries.

## Methods

In what follows, I provide an overview of the modelling framework, followed by detailed information on the study species, and of the demographic and climate modelling approaches. To assess the impact of managed culls and extreme climate conditions on the population dynamics of a marine mammal on the west coast of South America, I parameterised a stochastic matrix population model (MPM, hereafter) for the South American sea lion (*Otaria flavescens*) based on species-specific vital rate data, and vital rates imputed using phylogenetic comparative methods from other closely-related pinniped species. The resulting variance in these imputed vital rate estimates informed a distribution of matrix population models. Using a population vector of stage-specific population estimates, I then projected the population under different management and environmental scenarios. To assess the impacts of culls and extreme climate events on *O. flavescens* population viability, I conducted eight separate analyses of the population dynamics of *O. flavescens* over a 30-year time horizon. First, I assessed a base case condition (I), without managed culls and in the absence of extreme climate impacts. Then, I assessed the impact of three population cull scenarios: (II) 15% of adult females in year one, (III) 10% of adult females every year, and (IV) 30% of adult females every five years. Next, I assessed the impact of extreme climate events (V), represented by extreme El Niño conditions whose frequency I modelled using a Markovian transition matrix and whose impacts on vital rates were informed by observations from previous extreme El Niño years (Sielfeld and Guzmán, 2002; Soto et al., 2004). Finally, I assessed the combined impacts of the same three cull scenarios in combination with extreme climate events (VI-VIII) (Figure 1). Below, I describe each of these steps in more detail.

**Figure 1.**
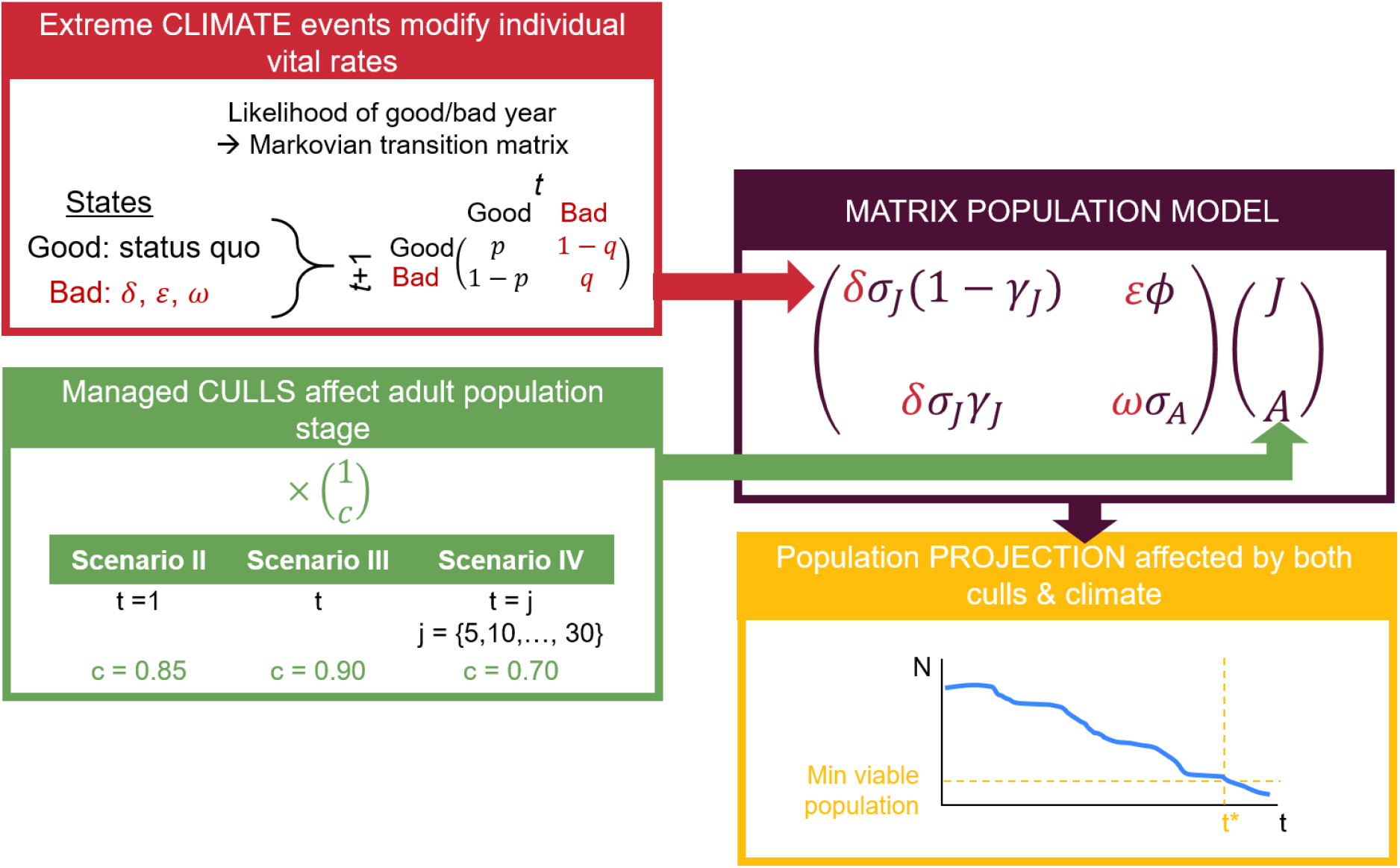
Overview of methodological pipeline to project the population dynamics of the South American sea lion (*Otaria flavescens*) using a matrix population model (dark purple panel) under the separate and combined impacts of managed culls (green panel) and extreme climate impacts (red panel). The vital rates of the matrix population model are: *σ* = survival; *Φ* = fecundity; and *γ* = maturation, representing juvenile females and adult females. Stage-specific managed culls remove adults from the population with different intensities (*c* = proportion of adult population removed) and frequencies (*t* = year). Extreme climate events, here extreme El Niño events, will alter vital rates by the terms *δ* (impacts on juvenile survival), *ε* (impact on fecundity), and *ω* (impact on adult survival). The frequency with which extreme climate events will occur is controlled through a Markovian transition matrix, where *p* = probability of climate remaining good, and *q* = probability of climate remaining bad. The population (*N*) of *O. flavescens* is projected over time (yellow panel) and assessed with relation to the year (*t*\*) in which the population drops below a minimum viable population size (dashed yellow lines).

### Study species

The South American sea lion (*O. flavescens*) is found along the South American coast, from Peru to Brazil, with a global population of ~400,000 individuals. Of these, 50% are found in Chile (Dans et al., 2004; Oliva et al., 2020). The species has been described as having plastic trophic habits, with diet determined by local prey abundance (Hückstädt and Antezana, 2006). In Chile, prey species include anchovy (*Engraulis ringens*), elephant fish (*Callorhynchus callorhynchus*), and hake (*Merluccius gayi*) (Hückstädt and Antezana, 2006). *O. flavescens* is one of the largest and most sexually dimorphic otariids (family Otariidae, describing pinnipeds with ears) (Cappozzo and Perrin, 2009), with adult males reaching 3m in length and 300-350kg, and adult females reaching 2m in length and up to 150kg (Cappozzo and Perrin, 2009). The species is polygynous, with males attempting to mate with as many females as possible (Cappozzo and Perrin, 2009). In Chile, the ratio of adult males to females in 2019 was ~1:11 (Oliva et al., 2020). Adult females typically produce one pup (Soto et al., 2004) each year, and the sex ratio at birth is 1:1 (Cappozzo and Perrin, 2009). In this analysis, I adopted a simplified female-only life cycle with pup and juvenile stages collapsed into a single ‘juvenile’ stage. This approach allowed me to better accommodate the emergent uncertainty from the available field data than more complex life-cycle models.

#### Vital rate estimation

I combined species-specific and imputed vital rates to parameterise stochastic MPMs for *O. flavescens*. Some information on vital rates for *O. flavescens* was available from Sepúlveda et al. (2006). Specifically, these authors estimated juvenile maturation to adult and adult fecundity for *O. flavescens* populations in the central regions of Chile. Other vital rates needed to be imputed due to the lack of available field information. Following recent findings regarding the robustness of phylogenetic methods to impute vital rates and life history traits (James et al., 2020; Johnson et al., 2021; Penone et al., 2014), I imputed juvenile survival and maturation and adult female survival using the Rphylopars package (Goolsby et al., 2017) in R (R Core Team, 2021). This method imputes missing data across a set of phylogenetically-related species (11 pinniped species in this case, including the target species) using a phylogeny and sparse trait matrix to simultaneously estimate phylogenetic and phenotypic trait covariance. I sourced vital rates for 10 pinniped species sourced from the COMADRE Animal Matrix Database v.4.20.11.0 (Jones et al., 2021). COMADRE houses 3,321 MPMs for 415 animals worldwide. In the case of pinnipeds, six of the 10 species contained >1 MPM, while four species were represented by only 1 MPM. Available matrices described unmanipulated populations (*i.e*., no experimental treatments).

For the variable number of stages in the 11 pinniped species to be consistent with my 2-stage life cycle (Fig. 2A), I collapsed available matrices to 2 × 2 stages (juveniles and adults) following methods by Salguero-Gomez and Plotkin (2010), using the R package Rage (Jones et al., 2021). Also using the R package Rage, I estimated stage-specific vital rates (mean and standard deviation) for each species. For the six pinniped species with multiple MPM available, I sampled 1,000 estimates of each vital rate based on the previously estimated mean and standard deviation, drawing from a truncated normal distribution (0-1 for survival and maturation, and 0-infinity for fecundity). I sourced a phylogenetic tree for mammals from Vertlife (Upham et al., 2019), which I pruned to the 10 pinniped species with MPM data from COMADRE and *O. flavescens* using R package ape (Paradis and Schliep, 2019). Using Rphylopars (Goolsby et al., 2017), I imputed missing vital rates for *O. flavescens* for each combination of the 1,000 estimates of vital rates for females of related species. I then parameterised MPMs for each of these 1,000 estimated vital rate values.

**Figure 2.**
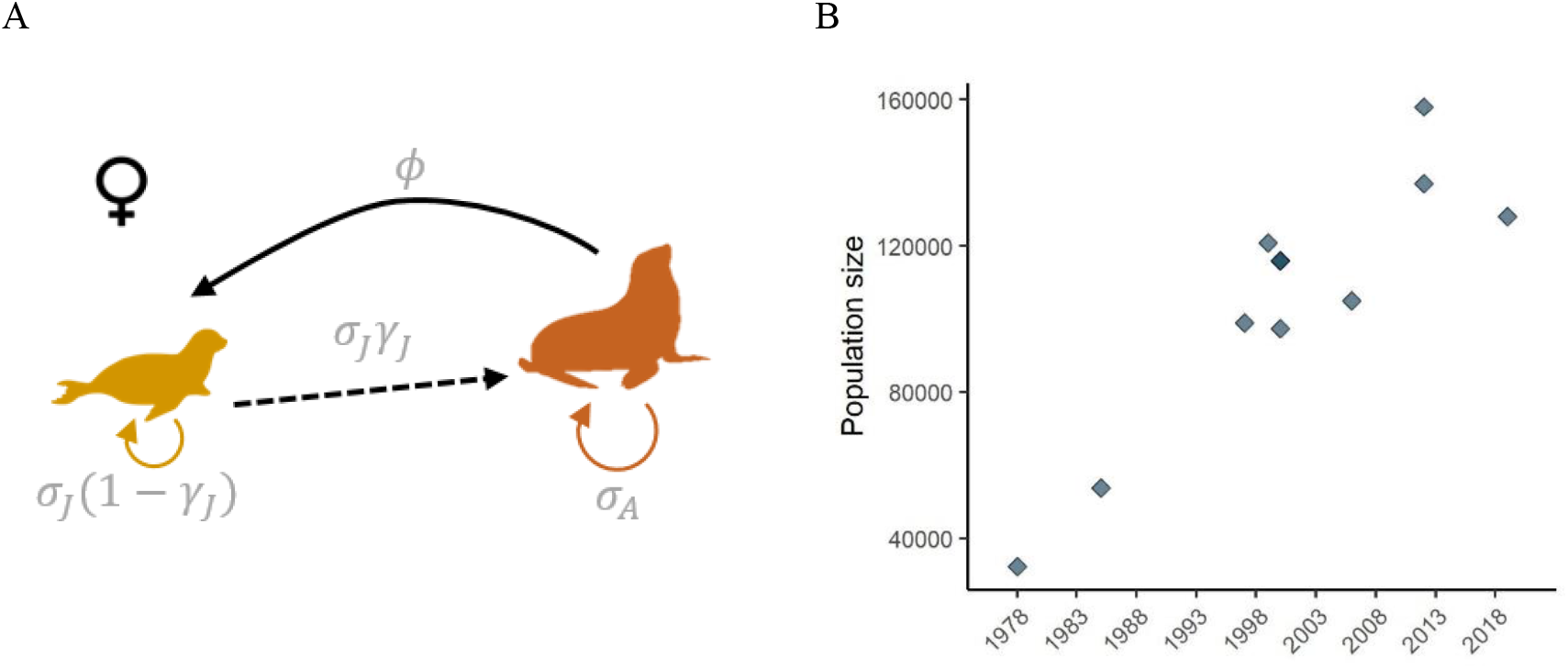
A. Life cycle of female *Otaria flavescens* (South American sea lion). Stages: J = juvenile female (< 4 years), A = adult female (≥ 4 years); Vital rates: *σ* = survival; *Φ* = fecundity; *γ* = maturation. B. Trend in abundance over time of the total *O. flavescens* population along the coast of Chile.

#### Population data

I combined a vector of stage-specific population data with the MPM to project the population through time. Trends in the total Chilean population of *O. flavescens* estimate are shown in Figure 2B. Stage-specific population estimates for *O. flavescens* were available for 2019 from Oliva et al. (2020). In this most recent census (2019) for Chile, there were an estimated 78,709 adult females, 7,126 juveniles, and 29,827 pups. Assuming a sex ratio of 1:1 in pups and juveniles, I calculated a combined pup and juvenile estimate for females (hereafter referred to simply as ‘juvenile’) of 18,477. These data provided a two-stage population vector for 2019 of 78,709 adult females and 18,477 juvenile females for the entire Chilean population (Figure 2).

To understand the base case dynamics of *O. flavescens* (scenario I), I projected the population of juvenile and adult females over 30 years (2019 to 2049) without any impacts of cull or extreme climate impacts. To do so, at time *t* + 1, I sampled a single MPM from the 1,000 estimated MPMs, and multiplied this matrix by the population vector in time *t*. I iterated this process 1,000 times.

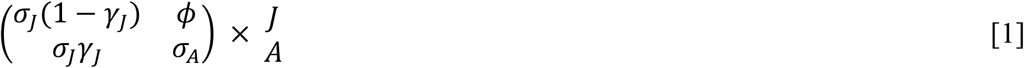

The top left hand term of the matrix describes the probability that a juvenile female (J) survives (*σ_j_*) but remains a juvenile in year *t*+1. The bottom left hand term of the matrix describes the probability that a juvenile female survives and matures (*γ_j_*) into an adult (A) in year *t*+1. The top right hand term of the matrix describes the number of juvenile females that an average adult female contributes to the population each year. Finally, the bottom right hand term of the matrix describes the probability an adult female survives (*σ_A_*) to year *t*+1. The two-element vector (*J*, *A*) describes the population size of juvenile females and adult females in year *t*.

To evaluate whether the population of *O. flavescens* will become functionally extinct during the modelled time horizon, I assessed whether and at what point in time the projected population drops below a minimum viable population (MVP, *sensu* Shaffer, 1981). I used a MVP value of 5,000 breeding adults, thought to be sufficient to ensure long-term population persistence and to avoid evolutionary decay across a broad range of taxonomic groups (Traill et al., 2010). Based on the sex ratio observed in the Chilean population (1:11.45 adult males to females), this criterion implies a MVP for adult females of 4,598 individuals. I quantified the time at quasi-extinction as the year of projection when >95% of my 1,000 simulated projected runs produced a population size (*N*) below this MVP value (Morris and Doak, 2002).

### Cull simulations

I assessed the impact of three different cull scenarios on the population of *O. flavescens* in Chile over 30 years. The three scenarios I considered were:

- Scenario II. Cull 15% of adult females in year 1 only;
- Scenario III. Cull 10% of adult females every year; or
- Scenario IV: Cull 30% of adult females every 5 years.

These cull levels mimic a range of harvest conditions previously considered in Chile (Sepúlveda et al., 2006). I simulated these culls on adult females rather than juveniles as culling in pinnipeds has historically targeted adult females due to their better accessibility relative to males or juveniles, particularly during the breeding season (summer) when adult females come to shore to pup. To estimate the impact of managed culls, I assessed the population dynamics of *O. flavescens* as described previously, but with the relevant proportion of the adult female population removed from the population vector at the appropriate time step, e.g., year 1 in scenario I, and every five years for scenario III.

### Climate conditions

In scenario V, I assessed the impact of extreme climate events on the population of *O. flavescens* in Chile. To model transitions between ‘good’ and ‘bad’ climate years, I used a Markovian transition matrix. Here, bad climate years described extreme El Niño events (Cai et al., 2014), events for which austral summer rainfall is greater than 5 mm per day. These extreme El Niño events, which included the 1982/83 and 1997/98 El Niño events, are characterised by exceptional warming, with sea surface temperatures exceeding 28°C extending into the eastern equatorial Pacific (Cai et al., 2014). These conditions lead to large decreases in prey availability (de Oliveira et al., 2012) and cause widespread environmental disruptions in the Pacific and beyond. Previous research in Peru, which neighbours Chile to the north, describes a 100% mortality of juveniles and 60% mortality of adults during the 1997/98 El Niño (Soto et al., 2004). Decreases in fecundity of 95% were also observed during this period (Soto et al., 2004). Sielfeld and Guzmán (2002) estimated similar levels of pup and juvenile mortality for populations in northern Chile. To represent these effects in my simulations, I modified each of the 1,000 combinations of vital rates previously estimated as below.

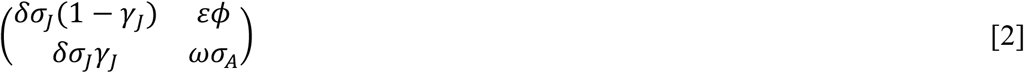

Where *δ* indicates the impact of extreme climate events on juvenile survival, *ε* indicates the impact on fecundity, and *ω* indicates the impact on adult survival. I parameterised these elements according to previous estimations as a 100%, 95%, and 60% decrease in the respective vital rates (Soto et al., 2004). This approach provided a distribution of matrix population models for ‘bad’ climate years. The population projection was as previously described, but this time, a Markovian transition matrix determined whether the population vector in *t* + 1 was multiplied by a matrix from the good or bad distribution.

El Niño events typically last 12-18 months (NOAA, n.d.), and the frequency of extreme El Niño events is predicted to increase under global warming (Cai et al., 2018). Relative to a ‘control’ period (1891-1990), when the frequency of extreme El Niño conditions was estimated at one event every 20 years, under current and future climate conditions (1991-2090), extreme events are predicted to occur once every ten years (Cai et al., 2014). I used this prediction to parameterise the Markovian transition matrix. When climate was good in time *t*, the probability of climate transitioning to bad in *t* + 1 was 0.1 (1 – p, Figure 1, red panel). When climate was bad in time *t*, the probability of climate transitioning to good in *t* + 1 was 0.5 (1 – q, Figure 1, red panel).

### Culls and climate

In scenarios VI-VIII, the previously described processes: culls and extreme climate conditions, were combined. I assessed the same three cull scenarios (II-IV), and in each of these scenarios I incorporated extreme climate conditions as previously described in scenario V.

### Sensitivity analysis

To identify how the population viability of *O. flavescens* is affected by extreme El Niño conditions, I assessed the sensitivity of results to different impacts of extreme El Niño conditions on vital rates. For each of the vital rates: (i) juvenile survival, (ii) juvenile maturation, (iv) adult survival, and (v) fecundity, I assessed a range of possible impacts varying from no impact, to a 100% decrease in each vital rate. I discretised this range of impacts into five levels: 0.0, 0.2, 0.4, 0.6, 0.8, and 1.0. For each of the 1,000 combinations of vital rates previously estimated, I modified the relevant vital rate by each level of the discretisation and conducted the population projection as previously described for the base case extreme climate scenario (scenario V).

## Results

The average MPM estimated for *O. flavescens* in Chile is:

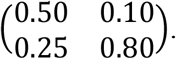

This MPM describes a juvenile female as having a 50% chance of surviving and remaining a juvenile, and a 25% chance of surviving and becoming an adult. Adult females have an 80% chance of survival, and contribute 0.1 juvenile females to the population each year. Based on this MPM, the average long term growth rate for the female *O. flavescens* population is 0.88 (95% confidence interval: 0.87, 0.88), which suggests that the population will decrease over time.

The matrix element with the largest impact on the per-capita population growth rate (*λ*) at equilibrium is adult survival (the bottom right element in the matrix below). A small unit change in this matrix element will increase *λ* by 76%. By contrast, there is little impact on *λ* of perturbations to fecundity, or juvenile stasis—whether a juvenile survives and remains a juvenile—or juvenile survival and maturation to an adult. The elasticity of *λ* to small unit changes in each of the matrix elements is shown below:

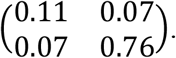

The Chilean population of *O. flavescens* is projected to decline over the next 30 years. Specifically, under the base case scenario (I), and based on the initial population vector of 78,709 adult females and 18,477 juvenile females, the population projection for *O. flavescens* in Chile describes a population decline over the next 30 years (Figure 3.I). This result is consistent with the estimated value of *λ*<1. Within this assessed time horizon (30 years) the population falls below the MVP size of 4,598 adult females (assessed as >95% of iterations meeting this criteria). This suggests that the population will not persist or avoid evolutionary decay within this time period.

**Figure 3.**
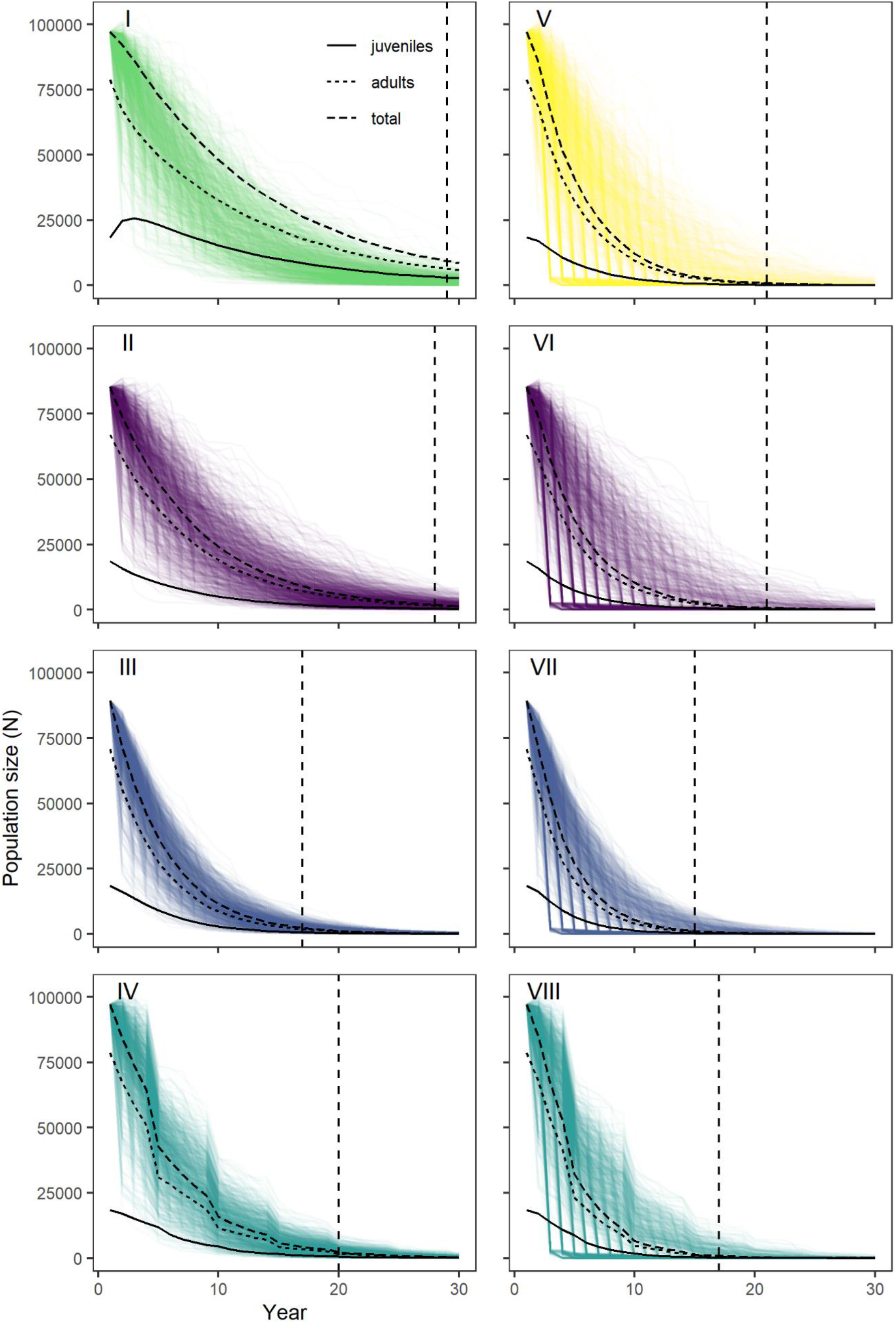
Population projections of combined adult and juvenile females of *O. flavescens* in Chile, where each line represents a model iteration (n = 1,000). Time horizon is 30 years with an initial population vector of 78,709 adult females and 18,477 juvenile females. Black vertical lines show stage-specific average values across all iterations. Horizontal dashed line indicates year where probability of all iterations dropping below the quasi-extinction threshold (4,598 adult females) is > 95%. I: Base case scenario. II: Cull 15% of adult females in year 1. III: Cull 10% of adult females every year. IV: Cull 30% of adult females every 5 years. V: Extreme climate impacts. VI: Cull 15% of adult females in year 1 and extreme climate impacts. VII: Cull 10% of adult females every year and extreme climate impacts. VIII: Cull 30% of adult females every 5 years and extreme climate impacts.

The *O. flavescens* population in Chile is projected to decrease below MVP under all of the assessed cull scenarios (scenarios II-IV, Figure 3). Specifically, this threshold is reached in year 28 when 15% of adult females are culled in year 1 only (scenario II), in year 17 when 10% of adult females are culled every year (scenario III), and in year 20 when 30% of adult females are culled every five years (scenario IV). Culling 10% of adult females each year therefore has the most extreme impact on the population dynamics of *O. flavescens* in Chile. By contrast, a population cull of 15% of adult females in year one (scenario II) has negligible impacts on the population relative to the base case scenario (scenario I). Under the projected frequencies of extreme climate events (scenario V), 34.3% of model iterations decline to <100 adult females within 10 years (Figure 3.V). However, the population is not assessed as quasi-extinct until year 21.

The combined impact of culls and extreme climate events is to speed up the population decline of *O. flavescens*. When the population of *O. flavescens* in subjected to culls and extreme climate events (scenarios VI-VIII), the population reaches quasi-extinction under all assessed cull regimes, and declines much faster than in cull scenarios without extreme events (Figure 3.VI-VIII). Under an initial cull of 15% of adult females in year one (scenario VI), extreme climate impacts lead to the population reaching quasi-extinction in year 21—7 years earlier than in the cull-only scenario (II). Quasi-extinction is hastened by 2 years in the case of an annual cull of 10% of adult females (to year 15, Figure 3.VII), and by 3 years in the case of a 30% cull of adult females every five years (year 17, Figure 3.VIII). An annual cull of 10% of adult females (scenario VII) remains the cull regime with the largest impact on population dynamics over the assessed time horizon when combined with extreme climate impacts. It should also be noted that the proportion of populations declining to very low numbers (< 100 adult females) increases in all cull scenarios with the combined impact of extreme climate events. Across scenarios VI-VIII, between 34.9%, and 36.1% of all iterations fall below 100 adult females within the first 10 years, compared to none in scenarios II-IV.

Based on these results, it is clear that the separate impacts of managed culls or extreme climate events would adversely affect the population of *O. flavescens* in Chile. In all assessed scenarios, the population declines below a MVP threshold over a 30 year time horizon when subject to culls (scenarios II-IV) or extreme climate events (scenario V). Under the combined impacts of managed culls and extreme climate events, the speed at which the population will become functionally extinct increases. Under all combined scenarios (VI-VIII) the population of *O. flavescens* in Chile falls below an MVP within 17-21 years.

In keeping with the analysis of the elasticity of the per-capita population growth rate (*λ*), a sensitivity analysis of the impact of extreme climate events on vital rates (Table 1) shows the greatest change in quasi-extinction occurs when adult survival is affected. Specifically, time to quasi-extinction is increased to 25 or 26 years when extreme climate events do not affect adult survival, or only reduce adult survival by 20%. Table 1 shows the different levels of extreme climate event impacts that were assessed. These range from no impact (0% decrease) to a 100% decrease in the relevant vital rate value. For vital rates other than adult survival, there is little difference in the speed with which populations decrease below the MVP threshold irrespective of the assumed impacts of extreme climate on each vital rate.

**Table 1.**
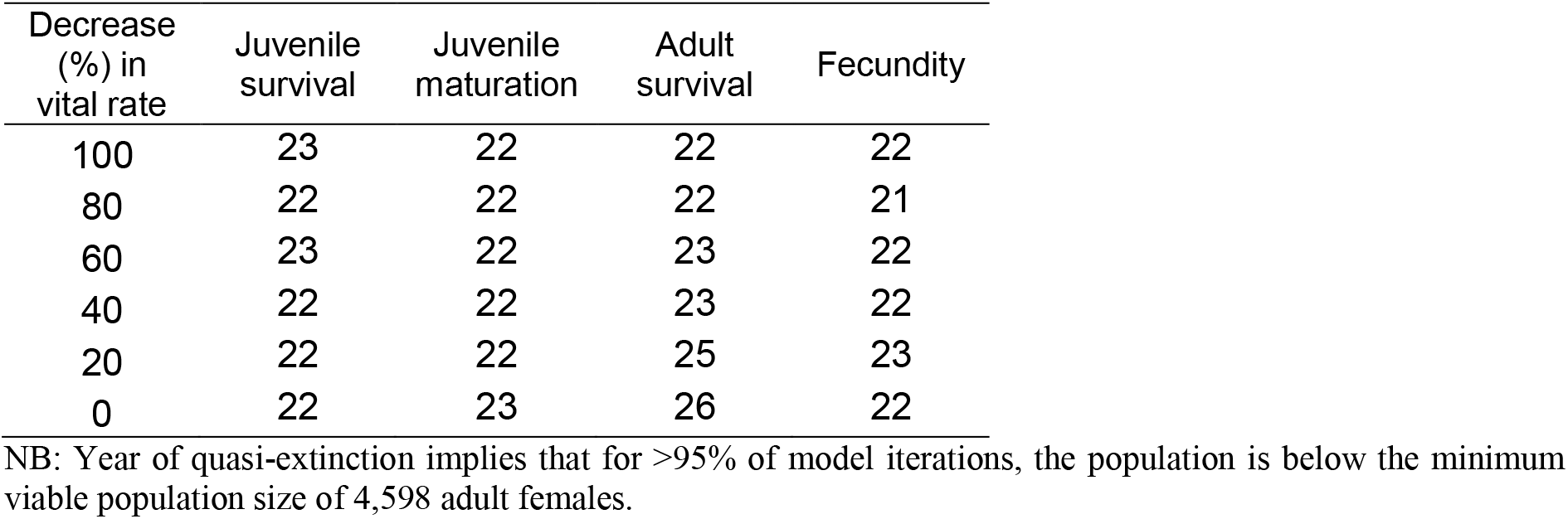
Time to quasi-extinction (years) of the Chilean population of the South American sea lion (*O. flavescens*) under different simulated impacts on vital rates of extreme El Niño events.

## Discussion

Using a stochastic matrix population model, I show that the Chilean population of the South American sea lion (*Otaria flavescens*) is likely to decline below a MVP threshold over the next 30 years. This demographic model estimates the long-term population growth rate of the Chilean population of this species at 0.88, indicating a significant population decline of 22% every year under current conditions. This key finding contrasts with the widespread perceptions of fishers in Chile that sea lion populations are too large and continue to grow (Davis et al., 2021). This dichotomy may be explained by a shifting baseline syndrome (Pauly, 1995). Although many sea lion populations around the world, and in Chile, have not yet recovered to historical population abundance (*i.e*., levels observed before widespread sealing in the 18^th^ to 19^th^ centuries) (Lotze et al., 2011; Magera et al., 2013), many present-day fisheries began operations in the years when sea lion populations were still small. Hence, current increases in these populations are assessed against the prevailing conditions when fishers began fishing (e.g., in the second half of the 21^st^ century) and thus are seen as ‘population explosions’ (Davis et al., 2021) instead of a recovery towards historical levels.

The speed of *O. flavescens* population decline will further increase under managed culls and/or expected frequencies of future extreme El Niño events. When managed culls and extreme El Niño impacts are combined, model simulations indicate a faster population decline. Based on these results, the Chilean population of *O. flavescens* is unlikely to support a managed cull—particularly given current expectations about the frequency of extreme climate events, including El Niño conditions. In keeping with these results, management efforts to reduce conflicts with fisheries should avoid managed culls—focusing on protecting and increasing populations rather than reducing them. Alternative conflict-management options may include capacity building schemes, financial compensation or growth in eco-tourism enterprises (Davis et al., 2021).

This analysis has demonstrated that extreme climate events will hasten the decline of *O. flavescens* below a MVP value. This finding indicates that the precautionary principle may need to be exercised when managing the population of *O. flavescens* on the west coast of South American—larger populations will be required to allow the population to buffer against extreme climate events. A similar conclusion was reached by de Oliveira et al. (2012) in an assessment of the effective population size of *O. flavescens* in Peru that explicitly considered the impacts of the mating system and demographic variations caused by the 1998-98 extreme El Niño event. In this work, the authors recommended a population of 7,715 individuals to ensure the population was large enough to avoid inbreeding and retain sufficient adaptive genetic variation to survive future El Niño events.

Model iterations show considerable variation in the assessed population projections through time, particularly in scenarios with low frequency shocks from culls or climate impacts (scenarios IV-VIII). These short-term fluctuations may be responsible for the broadly held perception that sea lion populations in Chile are rapidly increasing (Davis et al., 2021). These fluctuations can be explained by short-term, or transient, dynamics (Stott et al., 2010). Previous work has found that faster-growing populations tend to have relatively higher transient amplification (increase) or attenuation (decrease in population size) (Stott et al., 2010). In the current analysis, low asymptotic growth rates (0.88, 95% CI = 0.87-0.88) for female *O. flavescens* in Chile suggests this population has reduced capacity for short-term amplification or attenuation in population size. Transient amplification and attenuation have also been shown to be affected by matrix dimension (Stott et al., 2010)—this would suggest that additional complexity in the life cycle used to model *O. flavescens* (and hence in the matrix dimensions) would lead to increased short-term fluctuations in population size. Finally, Stott et al. (2010) also found that plant species with extreme life histories (e.g., monocarpic plants and trees) have greater potential for transient amplification and attenuation. The relevance of these findings for animal populations has yet to be tested, however, *O. flavescens* and other pinniped species tend towards the slow-end of the fast-slow life history continuum—suggesting they may have greater potential for short-term increases and decreases in population size.

If extreme climate events caused less adult mortality, it is likely that these events would have less impact on sea lion viability. A sensitivity analysis of extreme climate impacts on vital rates demonstrated that population viability was most sensitive to changes in the magnitude of extreme climate impacts on adult mortality, but relatively insensitive to changes in the magnitude of extreme climate impacts on other vital rates (juvenile survival and maturation and fecundity). It is worth noting that the impact of extreme El Niño events on vital rates was parameterised from northern populations (e.g., de Oliveira et al., 2012; Soto et al., 2004). It is unlikely that these impacts would be as severe for southern populations. If these impacts were weaker in southern colonies—particularly the impacts on adult mortality—then this will have positive implications for population viability. However, southerly populations may experience similar extreme population declines due to other stressors. For example, *O. flavescens* populations on the Argentinian coast decreased by 93% between 1938-1975 due to unknown causes (Gerber and Hilborn, 2001).

This analysis is one of only a few to incorporate uncertainty in vital rates into population projections (Paniw et al., 2017), and to my knowledge the first one to do so in pinnipeds. However, results would almost certainly be improved by further annual stage-based field data, which would allow species-specific vital rates to be estimated through inverse methods (Wielgus et al., 2008)—assuming that individual-based records were not feasible. In particular, more exact estimates of vital rates would allow a more complex life cycle to be assessed, including a two-sex model. As male animals interact more often with fisheries (Kauhala et al., 2012), a two-sex model would permit more nuanced modelling of cull or population control measures. In Chile, some level of illegal harvest of sea lions also occurs (Davis et al., 2021)— the impact of this mortality is not incorporated in the current analysis, as reliable data for these activities remains elusive. Data on these mortality levels would allow a better estimate of vital rates and of the long term population growth rate.

Conflict between marine mammals—including sea lions—and fisheries remains a highly contentious management issue (Davis et al., 2021; Ramos et al., 2020). This analysis has shown that this conflict will be best resolved independent of management culls. Perceptions of increases in sea lion populations may be an artefact of shifting baselines (Pauly, 1995), or the saliency of transient dynamics (Stott et al., 2010). Increases in the frequency of extreme climate events will leave sea lion populations vulnerable to extinction—requiring larger populations to remain viable over time. This precautionary, but necessary, approach is likely to exacerbate conflict with fisheries, highlighting an urgent need to develop and test alternative conflict-management solutions.

## Acknowledgements

This research was supported by the UKRI Internal Research England Global Challenges Research Support Fund [0008129] to K.J.D.

